# Kinetic dissection of pre-crRNA binding and processing by CRISPR-Cas12a

**DOI:** 10.1101/2023.07.25.550589

**Authors:** Selma Sinan, Nathan M. Appleby, Rick Russell

## Abstract

CRISPR-Cas12a binds and processes a single pre-crRNA during maturation, providing a simple tool for genome editing applications. Here, we constructed a kinetic and thermodynamic framework for pre-crRNA processing by Cas12a *in vitro*, and we measured the contributions of distinct regions of the pre-crRNA to this reaction. We find that the pre-crRNA binds rapidly and extraordinarily tightly to Cas12a (*K*_d_ = 0.6 pM), such that pre-crRNA binding is fully rate limiting for processing and therefore determines the specificity of Cas12a for different pre-crRNAs. The guide sequence contributes 10-fold to the affinities of both the precursor and mature forms of the crRNA, while deletion of an upstream sequence had no significant effect on affinity of the pre-crRNA. After processing, the mature crRNA remains very tightly bound to Cas12a, with a half-life of ∼1 day and a *K*_d_ value of 60 pM. Addition of a 5’-phosphoryl group, which is normally lost during the processing reaction as the scissile phosphate, tightens binding of the mature crRNA by ∼10-fold by accelerating binding and slowing dissociation. Using a direct competition assay, we found that pre-crRNA binding specificity is robust to other changes in RNA sequence, including tested changes in the guide sequence, addition of a 3’ extension, and secondary structure within the guide region. Together our results provide a quantitative framework for pre-crRNA binding and processing by Cas12a and suggest strategies for optimizing crRNA design in some genome editing applications.

---

Class II CRISPR-Cas endonucleases have revolutionized genome editing and a broad range of research applications because of their ability to selectively target DNA sequences through complementarity with their crRNA (Wang and Doudna, 2023). Of the class II enzymes, Cas12a is thought to have particularly strong potential because it targets linear or supercoiled DNA with very high specificity and is extraordinarily simple, processing its own pre-crRNA to a mature crRNA form by endonucleolytic cleavage to generate the 5’ end of the mature form (Fonfara *et al*., 2016, Swarts *et al*., 2017, Zetsche *et al*., 2017, Strohkendl *et al*., 2018, van Aelst *et al*., 2019, Nguyen *et al*., 2022) (Fig. 1A). This self-processing enables a simple strategy for multiplexing in genome applications, with multiple pre-crRNAs produced in a single transcript for processing by Cas12a (Zetsche *et al*., 2017). In addition to genome editing in cells, Cas12a has been used for applications including base editing and trans-gene integration (Kleinstiver *et al*., 2019, Mohr *et al*., 2023).

**Figure 1.**
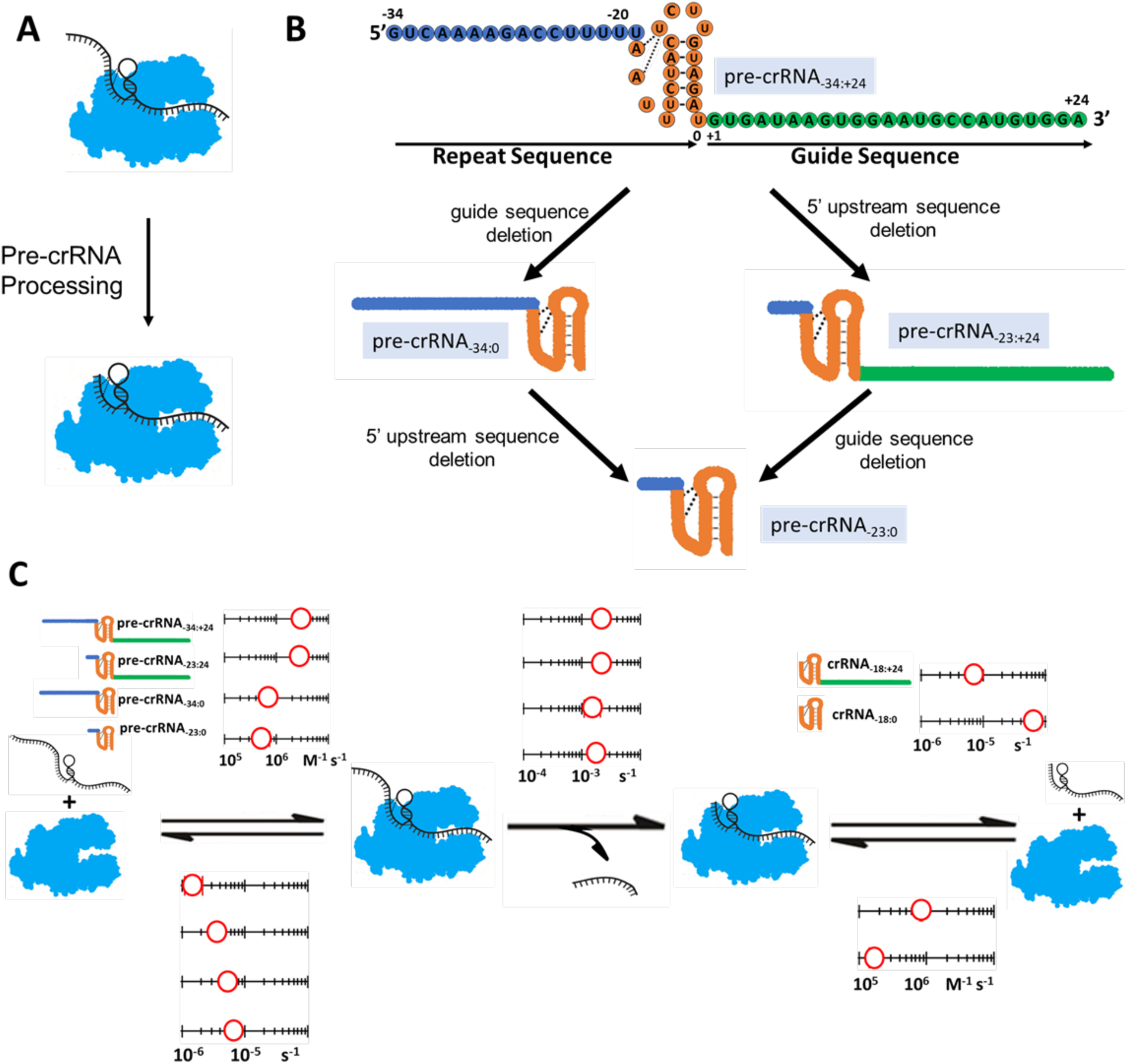
Kinetic framework for full-length and truncated pre-crRNA variants. (A) Processing reaction of pre-crRNA by Cas12a. (B) Sequence of the full-length pre-crRNA (pre-crRNA_–34:+24_), with the nucleotide numbering convention from (Swarts *et al*., 2017). The repeat sequence is divided conceptually into a leader sequence (blue) and pseudoknot (orange). The guide sequence is green. Deletion constructs used herein are schematized below the pre-crRNA sequence. (C) Binding, dissociation, and cleavage rate constants of the full-length pre-crRNA and the truncated forms tested herein. Binding and dissociation rate constants for the corresponding mature crRNA products are shown at the right. Uncertainties were typically 10-30% and are listed in Table S1.

The pre-crRNA can be divided into three major regions (Fig. 1B). At the 5’ end there is a leader sequence that is removed by endonucleolytic cleavage by Cas12a and is not part of the functional crRNA. Downstream of the leader is the ‘repeat’ sequence, which forms a pseudoknot structure and is critical for stable Cas12a binding (Fonfara *et al*., 2016). Following the repeat sequence is the guide sequence, known historically as the ‘spacer’, which forms an approximately 20-bp R-loop with the target strand DNA and is therefore responsible for much of the specificity in DNA targeting. In addition, there can be a fourth region comprised of any additional nucleotides downstream of the guide sequence, which are not removed by Cas12a and have been suggested to augment its function in certain cases (Bin Moon *et al*., 2018).

Analyses of the key features of the pre-crRNA for binding and processing have focused primarily on the pseudoknot region. When the RNA binds Cas12a to form a RNA-protein complex (RNP), the pseudoknot binds into a cleft between domains of Cas12a and is further stabilized by coordinating two hydrated Mg^2+^ ions (Swarts *et al*., 2017). Processing of the pre-crRNA occurs just upstream of the pseudoknot sequence and depends strongly on sequence and structural features of the pseudoknot (Fonfara *et al*., 2016). The chemical step does not itself require Mg^2+^ and is carried out by attack of the 2’-OH of nucleotide U(– 19) on the adjacent phosphoryl group, resulting in formation of a 2’, 3’-cyclic phosphate on the 5’ leader, which is released, and the mature crRNA, which remains tightly bound (Swarts *et al*., 2017). Although the affinity of the pre-crRNA has not been measured previously, it is known that binding of the processed crRNA is quite tight, with reported *K*_d_ values in the low nM range (Dong *et al*., 2016, Fonfara *et al*., 2016).

Because the functional properties of mature Cas12a RNP are largely controlled by the bound crRNA, there have been extensive efforts to engineer the crRNA for enhanced target specificity and other desirable properties. Modifications have included additional nucleotides at the 5’ end (Park *et al*., 2018) or the 3’ end (Bin Moon *et al*., 2018, Wu *et al*., 2018), splitting the crRNA into two RNA molecules (Jedrzejczyk *et al*., 2022, Shebanova *et al*., 2022), substituting segments of the crRNA with DNA nucleotides (Kim *et al*., 2020, Kim *et al*., 2022), and designing structures to compete against non-native structure involving the pseudoknot region (Kim *et al*., 2020, Kim *et al*., 2022). In spite of the utility of these efforts and some successes in cell-based assays, it can be difficult to generalize the effects or understand their molecular origin, and it is typically not known whether the functional effects reflect changes only in the properties of the Cas12a RNP itself or also reflect alterations in the interactions with other cellular components.

To further our mechanistic understanding of this enzyme, here we constructed a kinetic and thermodynamic framework for pre-crRNA recognition and processing by Cas12a. We found that binding of the pre-crRNA is extraordinarily tight (0.6 pM) and is fully rate-limiting for cleavage under subsaturating conditions. The 5’ leader sequence makes little or no contribution to pre-crRNA affinity, whereas the guide region contributes approximately 10-fold without a strong preference for a purine- or pyrimidine-rich sequence. The mature crRNA is also very stable, with a half-life of 26 hr despite an affinity that is 100-fold lower than that of the pre-crRNA. Interestingly, 5’ phosphorylation of the mature crRNA restores 10-fold of this lost affinity, indicating that applications requiring tight, fast binding of crRNA may benefit from this modification.

## RESULTS

### Kinetic and thermodynamic framework for pre-crRNA binding and processing

To probe the kinetic mechanism of pre-crRNA recognition and processing by Cas12a and to provide a foundation for experiments using mutant pre-crRNAs, we established a kinetic framework for pre-crRNA binding and processing by AsCas12a. We started by using a radiolabeled ‘full-length’ pre-crRNA (pre-crRNA_-34:+24_) consisting of a 35-nucleotide repeat sequence, including the pseudoknot structure, and a 24-nt guide sequence (Fig. 1B, top; Fig. S1). For each reaction step, we used polyacrylamide gel electrophoresis (PAGE) to separate the starting material from the ‘product’ of the reaction step being measured.

We first performed single-turnover pre-crRNA processing reactions under conditions of saturating or subsaturating Cas12a to measure the maximal rate constant (*k*_max_) and second-order rate constant (*k*_cat_/*K*_M_), respectively. Thus, we measured the time dependence of formation of the shorter mature crRNA by denaturing PAGE. With saturating Cas12a, the observed rate constant was 2.3 (±0.3) × 10^-3^ s^-1^ (Fig. 2A; results summarized in Fig. 1C and Table S1), which reflects the rate constant for pre-crRNA cleavage from the bound complex. With subsaturating Cas12a concentrations, the dependence of the observed rate constant on concentration gave a *k*_cat_/*K*_M_ value of 2.9 (±0.2) × 10^6^ M^-1^ s^-1^ (Fig. 2A). Experiments described below indicate that this value reflects the rate constant for pre-crRNA binding to Cas12a (*k*_on_).

**Figure 2.**
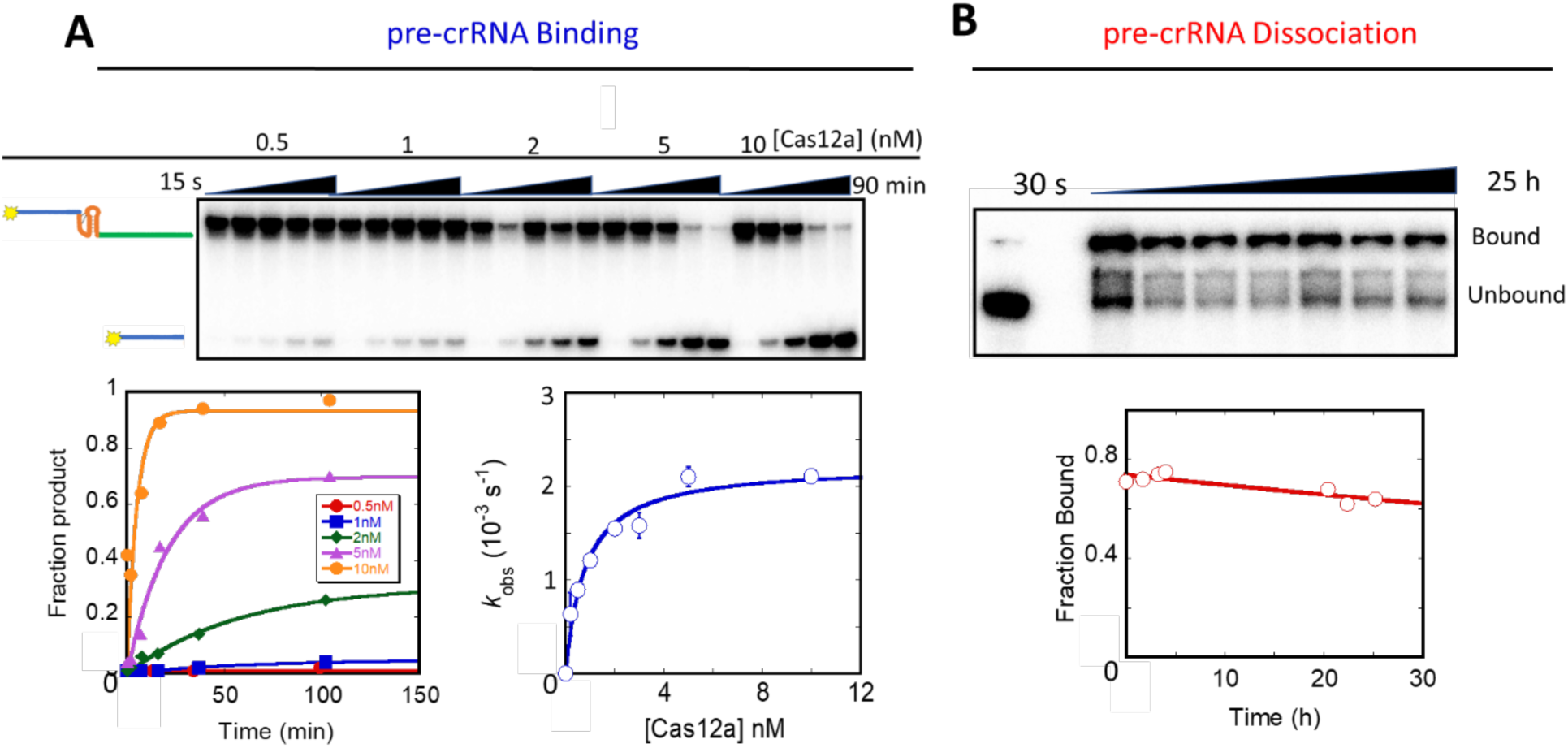
Cas12a binding and dissociation of pre-crRNA_-34:+24_. (A) Binding and pre-crRNA cleavage kinetics. The top image shows a representative gel, with the indicated Cas12a concentrations, trace radiolabeled pre-crRNA-34:+24, and time points from 30 s to 90 min. Data from the gel are shown as time courses (lower left), and observed rate constants from these and additional experiments are plotted against Cas12a concentration to give the second-order rate constant and maximal first-order rate constant indicated (lower right). Data are shown as the average and SEM of 3 independent measurements. (B) Dissociation kinetics. Trace radiolabeled pre-crRNA_-34:+24_ was bound to 5 nM Cas12a and chased off by addition of 50 nM unlabeled pre-crRNA_-34:+24_. The additional bands visible just above the free substrate in the gel image (top) were consistently observed in the presence of the chase oligonucleotide and probably reflect fortuitous interactions of two molecules of pre-crRNA_-34:+24_. The plot (bottom) shows results from the gel image.

Next we measured the dissociation rate constant of pre-crRNA from Cas12a by using a pulse-chase procedure, with bound and free pre-crRNA resolved by native PAGE (Fig. 2B). For these experiments, the pre-crRNA included a deoxyuridine at position −19 to prevent its cleavage by Cas12a (Swarts *et al*., 2017) (Fig. S2). Dissociation was very slow, giving a *k*_off_ value of 1.6 (±0.5) × 10^-6^ s^-1^ (Fig. 2B). This rate constant is 1000-fold lower than that for RNA processing by Cas12a, indicating that RNA binding is functionally irreversible and therefore that the *k*_cat_/*K*_M_ value measured above reflects the rate constant for pre-crRNA binding, *k*_on_. From the *k*_on_ and *k*_off_ values, we calculated an equilibrium constant value (*K*_d_) for pre-crRNA binding of 0.56 (±0.18) pM (*k*_off_/*k*_on_). This value reflects much tighter binding than the low nM value that was determined previously for binding of the mature crRNA by the related LbCas12a (Dong *et al*., 2016).

### Mg^2+^ ion accelerates processing and slows dissociation of pre-crRNA

Although early work indicated that divalent cations promote pre-crRNA processing by FnCas12a (Fonfara *et al*., 2016), later work showed that Mg^2+^ is not absolutely required for processing by LbCas12a, although Mg^2+^ or other divalent cations increased the extent of RNA cleavage (Swarts *et al*., 2017). To more fully understand how Mg^2+^ participates in pre-crRNA processing, we determined the effects of Mg^2+^ concentration on AsCas12a binding and processing of the full-length pre-crRNA (pre-crRNA_-34:+24_). We found that the cleavage rate constant was unaffected by Mg^2+^ concentration from 1 mM to 10 mM, and it decreased by only 5-fold in the absence of Mg^2+^ (Fig. S3). These results support the conclusion that pre-crRNA cleavage proceeds through nucleophilic attack by the 2’-hydroxyl group of the upstream ribonucleotide on the scissile phosphate (Swarts *et al*., 2017), and they indicate that one or more tightly bound Mg^2+^ ions play a minor but detectable role in the catalytic reaction, perhaps by binding and stabilizing the pseudoknot structure (Swarts *et al*., 2017) or by contributing to positioning of the substrate in the nuclease active site.

We also used native PAGE to measure the pre-crRNA binding and dissociation rate constants in the absence of Mg^2+^. The binding rate constant was minimally impacted by removing Mg^2+^ (6.0 (±0.3) × 10^6^ M^-1^ s^-1^), but dissociation was accelerated 19-fold to 3.2 (±1.5) × 10^-5^ s^-1^ (Fig. S4). Thus, the *K*_d_ value is 5.2 (±2.5) pM in the absence of Mg^2+^, 10-fold weaker than in its presence. This result is consistent with previous findings that Mg^2+^ promotes ordering of the crRNA pseudoknot and increases the RNA binding affinity of FnCas12a (Dong *et al*., 2016, Fonfara *et al*., 2016). Nevertheless, pre-crRNA binding by Cas12a remains very tight, in the low pM range, in the absence of Mg^2+^.

### Kinetics and thermodynamics of the Cas12a complex with mature crRNA

To probe further the interactions of Cas12a with crRNA and to aid in genome engineering applications, we determined rate and equilibrium constants for interaction of Cas12a with the mature, processed crRNA (crRNA_–18:+24_). We used native PAGE to measure the binding and dissociation rate constants of the mature crRNA, with the crRNA labeled at either the 3’ or 5’ end (Fig. 3). For the 3’-labeled crRNA, pulse-chase experiments analogous to those above gave a *k*_on_ value of 1.31 (±0.08) × 10^5^ M^-1^ s^-1^ and a *k*_off_ value of 7.4 (±0.9) × 10^-6^ s^-1^. These values gave an equilibrium constant of 56 (±8) pM and a half-life of 26 hr for the bound complex, indicating that the mature crRNA continues to bind very tightly to Cas12a and does not readily dissociate.

**Figure 3.**
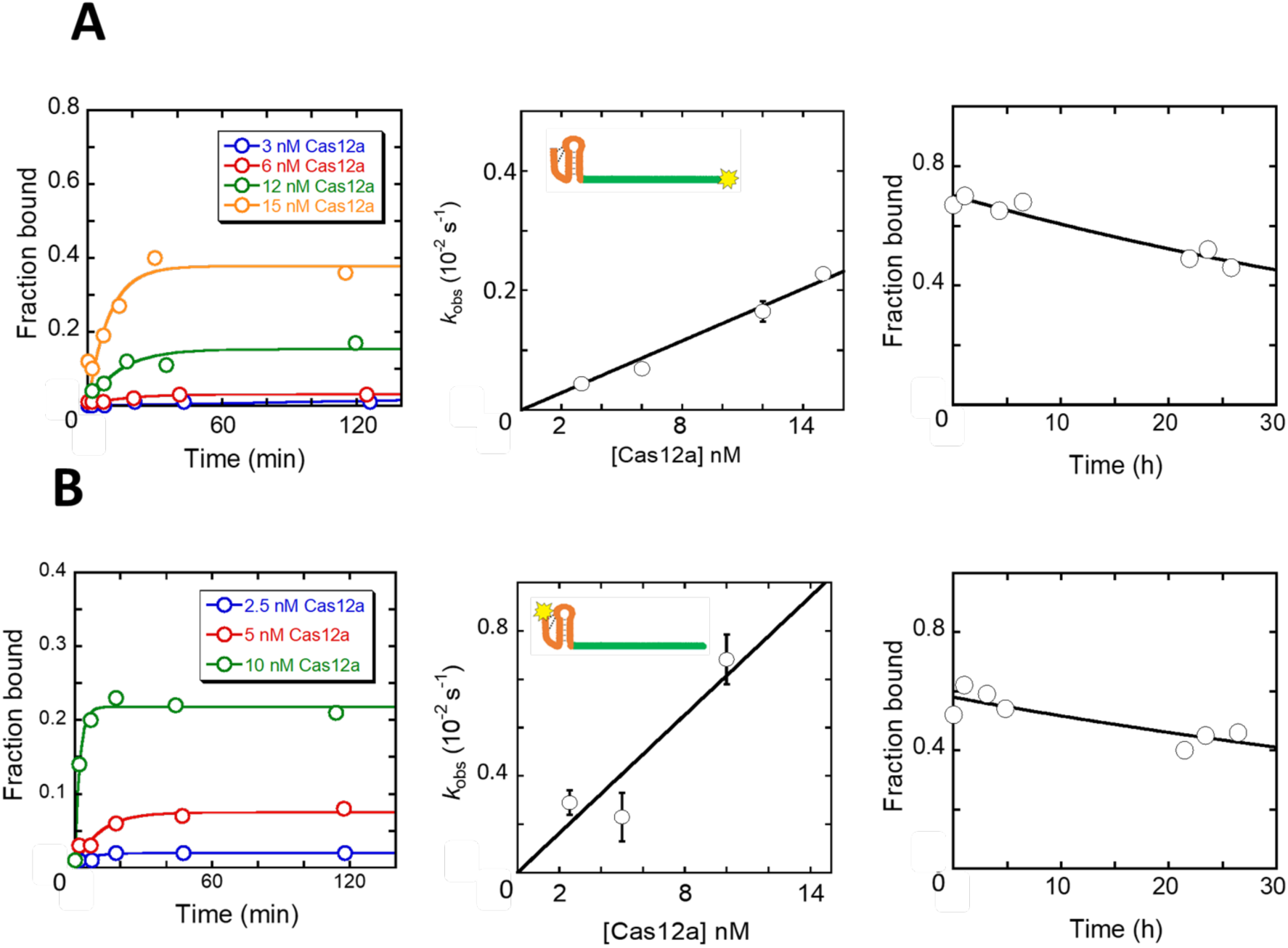
Binding and dissociation kinetics for mature crRNA. (A) Binding (left and center) and dissociation (right) kinetics of the mature crRNA_–18:+24_ with a 3’ label. In the center plot, binding data are shown as the average and SEM from at least 3 independent measurements. (B) Binding (left and center) and dissociation (right) kinetics of the mature crRNA_–18:+24_ with a 5’ label. In the center plot, binding data are shown as the average and SEM from at least 3 independent measurements.

For the 5’-labeled crRNA product, where the label is present on a phosphoryl group that mimics the scissile phosphoryl group in the processing reaction, we found that binding of this product was 6-fold faster than for the 3’-labeled product (8.3 (±0.3) × 10^5^ M^-1^ s^-1^) and dissociation was somewhat slower (4.5 (±0.03) × 10^-6^ s^-1^). Thus, the equilibrium constant was 5.4 (±0.2) pM for the 5’-labeled crRNA, ∼10-fold tighter binding than observed for the 3’-labeled RNA.

We hypothesized that the tighter binding of the 5’-labeled crRNA might reflect favorable contacts made by the phosphate bound at the active site. To test this hypothesis, we measured the lifetime of the Cas12a complex by performing multiple-turnover processing reactions, with a small excess of pre-crRNA over Cas12a (Fig. S5). With either 5’-labeled or 3’-labeled pre-crRNA, we observed a relatively rapid burst of product formation and then a slow phase of further product formation, which we infer reflects the slow release of the product crRNA. The rate constant for this slow phase was 2.2 (±0.3) × 10^-5^ s^-1^ for the 5’-labeled substrate. For this substrate, the label dissociates rapidly with the 5’ leader sequence after pre-crRNA cleavage, and thus the crRNA product whose rate is followed is unlabeled. We obtained a similar rate constant for a 3’ labeled substrate (1.1 (±0.1) × 10^-5^ s^-1^), and this value was also similar to the dissociation rate constant for this product measured above using pulse-chase procedures (7.4 (±0.9) × 10^-6^ s^-1^). Thus, the additional nucleotide that is added along the 3’ label contributes little to the lifetime of the complex with the crRNA (≤2-fold).

Together, the results indicate an equilibrium constant of 56 (±8) pM for the mature crRNA, approximately 100-fold weaker than binding of the pre-crRNA substrate but still extraordinarily tight binding for an RNA-protein complex and considerably tighter than the previously reported value of 3 nM for LbCas12a (Dong *et al*., 2016). Although the two Cas12a proteins may differ in RNA binding affinity, it is likely that the previous measurement underestimated the affinity because the incubation time used, 15 min, was likely insufficient to reach equilibrium (Bisaria *et al*., 2017). In addition, we find that the presence of a 5’-phosphoryl group contributes approximately 10-fold to equilibrium binding of the mature crRNA, reducing the *K*_d_ value to ∼5 pM. While the 5’ end is not phosphorylated in nature in the mature crRNA due to the processing mechanism, it may be useful to include this group artificially for certain gene editing applications (see Discussion).

### Dissection of the regions within pre-crRNA

An early study established the importance of the pseudoknot structure for pre-crRNA processing (Fonfara *et al*., 2016), but there is little knowledge of how other regions of the pre-crRNA contribute to assembly and processing by Cas12a. Therefore, we generated pre-crRNA variants with deletions of the 5’ upstream sequence (pre-crRNA_-22:+24_), the guide region (pre-crRNA_-34:0_), or both regions (pre-crRNA_-23:0_) (see Fig. 1).

We first measured pre-crRNA cleavage at saturating or near-saturating concentrations and found that none of these deletions gave large effects on the rate constant for pre-crRNA cleavage after assembly, as all three truncated versions gave the same rate constant (within 2-fold) as the full-length pre-crRNA_–35:+24_ (Fig. 4A; see also Fig. 1C). Next, from the dependence of the observe rate constant on subsaturating Cas12a concentrations, we found that deletion of the guide region slowed Cas12a binding by 2-4 fold (compare pre-crRNA_-34:0_ with pre-crRNA_–34:+24_; also compare pre-crRNA_-23:0_ with pre-crRNA_–23:+24_), while deletion of the upstream region had little or no effect on the binding rate constant (<2-fold). Deletion of the guide region also increased the dissociation rate constant by 2-3-fold. Together, these effects resulted in a calculated equilibrium constant (*K*_d_) of 5.0 (±1.3) pM for the double deletion construct, approximately 10-fold weaker binding than the full-length pre-crRNA. The variant pre-crRNA_–34:0_ gave approximately the same *K*_d_ value (7.2 (±1.4) pM), underscoring the conclusions that the guide region contributes an order of magnitude to binding affinity and the upstream sequence has little or no effect on binding.

**Figure 4.**
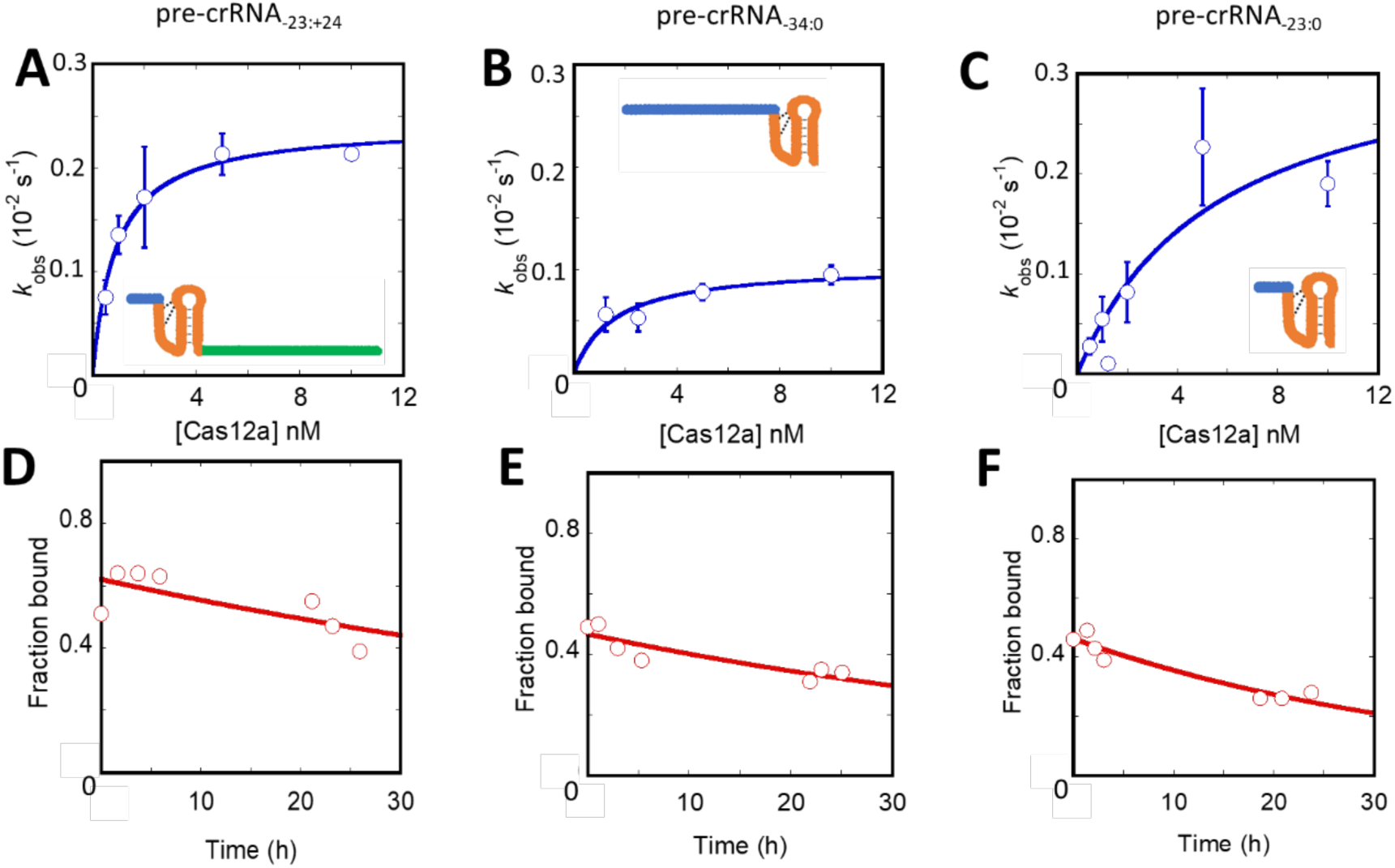
Binding and dissociation kinetics for truncated pre-crRNA variants. (A-C) Binding kinetics measurements for (A) pre-crRNA_-23:+24_ (B) pre-crRNA_-23:0_ (C) pre-crRNA_-34:0._ Data are shown as the average and SEM from at least four independent measurements. (D-F) Dissociation kinetics measurements for the Cas12a variants shown in panels A-C, respectively. Data are shown as the average and SEM from at least 3 independent measurements.

We also examined the effect of the guide region on binding of the mature crRNA, the product of Cas12a processing. By comparing the binding and dissociation rate constants for a truncated crRNA (crRNA_–18:0_) with those of the full-length crRNA_–18:+24_, we found that the guide region contributes 7-fold to the affinity of the mature crRNA product, via slowing dissociation by 9-fold while having little or no effect on the binding rate constant (Fig. S6). Thus, the guide region contributes approximately the same binding energy to the crRNA before and after its processing by Cas12a. For the crRNA, the effect of the guide region is almost entirely on the dissociation kinetics of the mature crRNA, whereas it impacts both the binding and dissociation kinetics for the pre-crRNA.

### Pre-crRNA binding specificity is robust to changes in length, sequence, and structural features of the guide region

We were particularly interested in the finding above that the presence of the guide region accelerates binding of the pre-crRNA, because it is this binding rate constant that controls the overall efficiency of Cas12a loading. Thus, the impacts of the guide region might influence the relative loading efficiencies of guide in multiplex genome editing applications as well as in biology. To further explore this possibility, we investigated how the guide sequence, length, and secondary structure impact the binding rate constant. To increase the sensitivity of detection for changes in the binding rate constant and to directly reflect the potential competitive environment in multiplex applications, we used a direct competition method in which each reaction included a reference pre-crRNA (either pre-crRNA_-34:+24_ or pre-crRNA_–22:+24_) and a given pre-crRNA variant, each in modest excess of Cas12a (Fig. 5). We expected that as Cas12a performed a fast, single round of pre-crRNA binding and processing, the amount of product produced from the pre-crRNA variant, relative to the amount of product from the reference pre-crRNA, would reflect its relative binding rate constant.

**Figure 5.**
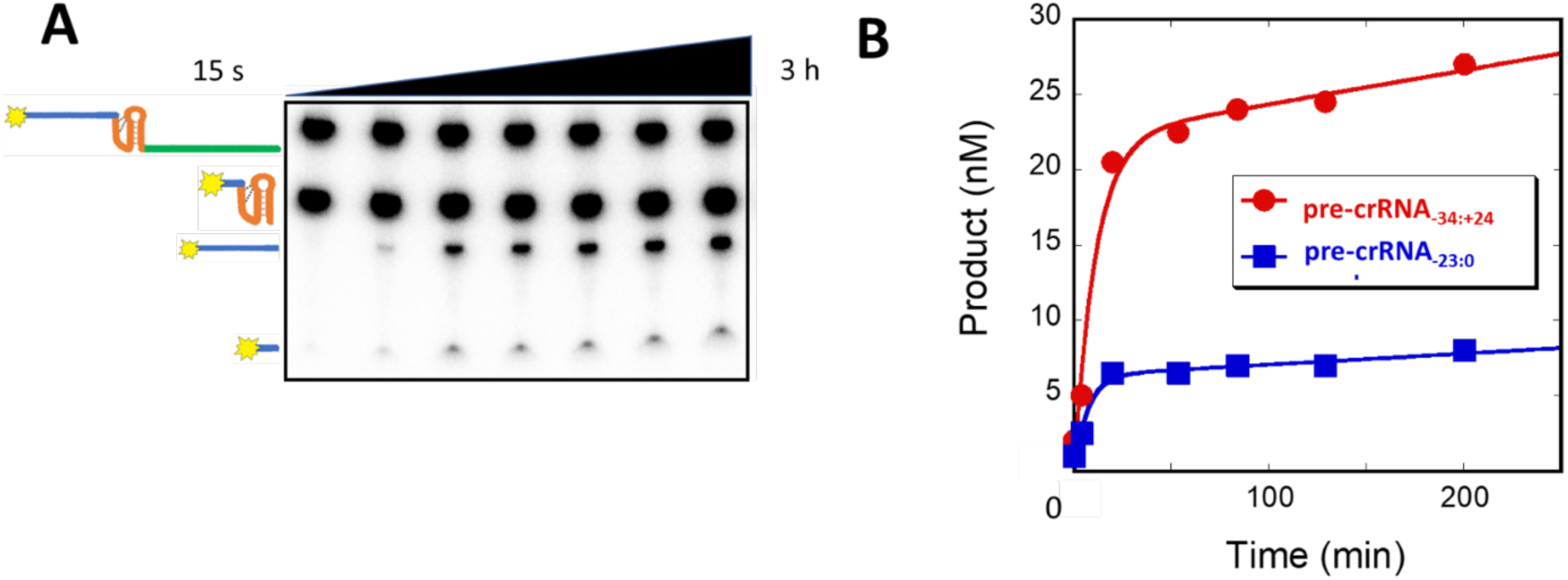
Competition experiment for Cas12a binding by pre-crRNA_-34:+24_ and pre-crRNA_-23:0_. (A) Denaturing gel image showing the substrates and products for both pre-crRNAs. The reaction included 50 nM of each pre-crRNA and 30 nM Cas12a. (B) Progress curves depicting cleavage of each pre-crRNA variant. Because Cas12a binding is rate-limiting for cleavage, the relative amplitudes of the ‘burst’ phase of product accumulation correspond to the relative rate constants for binding of the two pre-crRNA variants.

To ensure that the direct competition method gave results consistent with binding measurements, we first measured the effect of deleting the guide region. The amount of products, relative to the full-length pre-crRNA_-34:+24_ product, were 0.25 (± 0.08) for pre-crRNA_-22:0_ and 0.27 (± 0.12) for pre-crRNA_-34:0_ (Fig. 5; Fig. S7), in good agreement with the expected values of 0.21 (± 0.04) and 0.23 (± 0.02) from the binding rate constant measurements above (see Fig. 1). Also as expected, the pre-crRNA variant lacking the 5’-leader sequence competed equally with the full-length pre-crRNA, with a value of 1.0, consistent with the value of 0.87 from the rate constant measurements (Fig. 5).

Having established the competition method, we next investigated potential effects of the RNA sequence within the guide region by competing the pre-crRNA_-34:+24_ against variants with large changes in sequence; specifically the all-purine sequence A_20_ or the all-pyrimidine sequence U_20_. These variants gave relative binding rate constants of 0.63 ± 0.02 and 0.83 ± 0.07, respectively (Fig. 6; Fig. S7). These small effects in response to very large changes in sequence and base structure suggest that Cas12a is relatively insensitive to sequence changes in the guide region, despite the importance of its presence. We also investigated how the length of the guide impacts the Cas12a binding rate constant. A pre-crRNA with a 10-nt guide sequence gave a value of 0.69 ± 0.17, indicating minimal decrease in binding rate constant. Likewise, a pre-crRNA with eight uracil nucleotides (U_8_) appended to the 3’ end of the guide sequence gave a value of 0.69 ± 0.02. Because previous work indicated enhanced functionality in cell-based applications for a crRNA with U_8_ at the 3’ end of the guide region, we also measured dissociation of the processed form via multiple turnover experiments. We determined a rate constant of 2.16(±0.03) × 10^-5^ s^-1^, which is the same as that for pre-crRNA_-35:+24_, indicating that the lifetime of the crRNA-Cas12a complex is unaltered by the U_8_ addition. Together, our results suggest the absence of large effects of the sequence or length of the guide region on the Cas12a binding rate constant, although complete removal of the guide region reduces the binding rate constant ∼4-fold.

**Figure 6.**
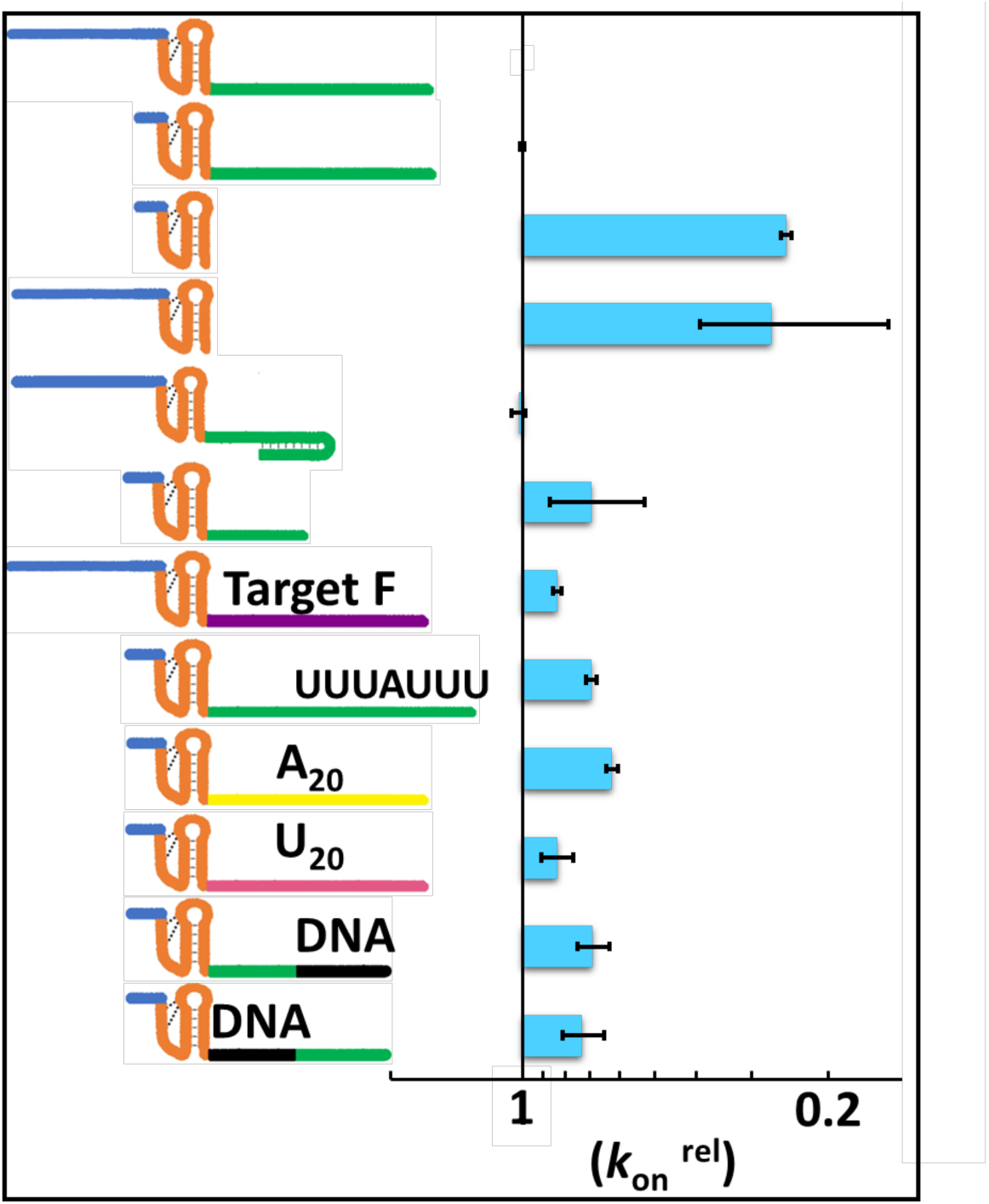
Relative Cas12a assembly rate constants measured by direct competition. Each pre-crRNA variant tested is shown in the cartoon at the left, and the corresponding bar at the right shows the rate constant relative to the reference pre-crRNA. The pre-crRNA at the top, pre-crRNA_-34:+24,_ is assigned a relative rate constant of 1 by definition. Binding of the variant shown immediately below it, pre-crRNA_-23:+24_, was measured and found to be equivalent in rate, as indicated by the absence of a bar to its right. All of the variants below were measured using one of these two pre-crRNAs as a reference, whichever one differed in length of the 5’ leader sequence from the variant being tested and therefore could be distinguished by the length of the labeled cleavage product.

Last, we measured the effects on binding rate constant of additional changes to the guide region with potential utility or consequences in cell-based applications. To probe the effect of secondary structure within the guide region, we measured competition of a pre-crRNA with a guide sequence that can form a 7-bp hairpin structure. This variant competed equally with the full-length pre-crRNA_-34:+24_, giving a ratio of 1.02 ± 0.037 (Figure 6). The Cas12a RNP loaded with this crRNA was strongly inhibited for DNA target recognition (Fig. S8), suggesting that the secondary structure of the crRNA remains formed in the mature Cas12a RNP. We also measured competition of pre-crRNA variants with chimeric, RNA-DNA guide regions in which DNA replaced nt 1-10 or 11-20 of the guide region. These variants gave competition values of 0.73 ± 0.06 and 0.69 ± 0.04, respectively, suggesting that DNA regions of the guide region have at most small effects on the binding rate constant. Because such chimeric guides are suggested to be useful for applications by increasing specificity of DNA recognition, we also measured the lifetimes of these chimeric crRNAs after processing, again using multiple turnover reactions. The dissociation rate constants were 4.3 (±0.9) × 10^-5^ s^-1^ and 2.6 (±1.3) × 10^-5^ s^-1^ for the variants with DNA in the proximal or distal positions, respectively. These values are at most modestly larger than the value of 2.2 (±0.3) × 10^-5^ s^-1^ measured for the standard crRNA, indicating that the substitution of up to ten DNA nucleotides does not compromise the lifetime of the crRNA-bound complex significantly.

## DISCUSSION

Our systematic kinetic analysis of the pre-crRNA binding and processing reaction provides insights into how Cas12a recognizes RNA and suggests strategies for improving Cas12a properties in some gene editing applications. We find that association of pre-crRNA with Cas12a is efficient and functionally irreversible, as processing occurs much faster than dissociation. The guide region of the pre-crRNA contributes significantly to binding affinity, both by increasing the pre-crRNA binding rate and decreasing the dissociation rate, while our work provides no evidence for effects of additional nucleotides upstream from the conserved pseudoknot.

While previous work established that the pseudoknot region is a key determinant of crRNA affinity for Cas12a, contributions of other regions of the crRNA had not been systematically analyzed. Our finding that deletion of the guide region results in a 10-fold decrease in affinity implies that there are contacts formed with Cas12a by this region. The crystal structure of FnCas12a revealed contacts between positions 1-5 of the crRNA backbone (*i.e.* the 5’ portion of the guide sequence) and amino acids in the WED and REC1 domains (Swarts *et al*., 2017). Interestingly, most of the 10-fold increase in affinity arises from an acceleration of binding, suggesting that contacts formed with the guide region are formed early in the binding process, prior to attainment of the rate-limiting transition state for binding. Recent structural and computational work indicates that pre-crRNA assembly can include a large-scale conformational change of Cas12a from an open to a closed complex (Jianwei *et al*., 2023, Sudhakar *et al*., 2023). This rearrangement may follow the establishment of most or all of the contacts between the pre-crRNA and Cas12a, completing the assembly process and limiting its overall rate.

After Cas12a cleaves the pre-crRNA to its mature crRNA form, binding of the crRNA remains strong and long-lived, properties that are presumably critical for the DNA-targeting function of the RNP. Nevertheless, the affinity of the mature crRNA is roughly 100-fold lower than that of the pre-crRNA, and further dissection of the energetic contributions revealed that 10-fold of this effect (i.e. half of the free energy difference) can be recovered by adding a 5’-phosphate group that mimics the scissile phosphate, which is released upon cleavage as part of the upstream leader sequence. Structural work showed that this phosphate group contacts a lysine residue at the active site, presumably to stabilize developing negative charge during the cleavage reaction (Swarts *et al*., 2017). Although the 5’ phosphate group is absent from the natural mature crRNA, applications of Cas12a that involve addition of pre-made, mature crRNA to Cas12a may benefit from the increased RNP lifetime that would be expected from its addition.

Overall, we found that the binding properties of the pre-crRNA to Cas12a are remarkably robust. Adding nucleotides to the 3’ end of the guide region or 5’ of the cleavage site, or replacing segments with DNA, did not significantly impact the kinetics or equilibrium binding properties. Likewise, varying the sequence of the guide region gave no detectable effect on Cas12a binding. These results suggest that findings of widely variable efficiency of different crRNAs for genome targeting or editing from such changes are unlikely to arise from differences in Cas12a binding. Even a pre-crRNA with a guide region that is predicted to form a stable hairpin binds Cas12a with the same binding rate constant, and the inability of the resulting RNP to efficiently target its cognate DNA indicates that the hairpin is indeed formed in the RNP. These results suggest that a crRNA with a guide region with self-complementarity would serve as an effective inhibitor, because it would bind readily to Cas12a, competing with other crRNAs, but would presumably be unable to recognize target DNA. This model is consistent with previous findings for Cas9 (Thyme *et al*., 2016) but runs counter to an earlier conclusion from cell-based fluorescence measurements suggesting that potential for base-pairing within the Cas12a guide region did not compromise crRNA function (Creutzburg *et al*., 2020). It is possible that RNA secondary structure changes in the guide region are facilitated by the intracellular environment, mitigating the inhibition of DNA targeting by internal crRNA secondary structure.

## METHODS

### Cas12a Expression and Purification

Cas12a was expressed as a fusion protein with a cleavable His6/Twin-Strep/SUMO N-terminal protein and purified as described (Strohkendl *et al*., 2018). After purification, small aliquots were flash-frozen in liquid nitrogen and stored at −80°C. Protein concentrations were determined by the Bradford assay (Bio-Rad) using bovine serum albumin as a standard.

### Measurement of pre-crRNA cleavage rates

Pre-crRNA was 5’-radiolabeled with [γ−^32^P] ATP (Perkin-Elmer) using T4 polynucleotide kinase (New England Biolabs). For some measurements, mature crRNAs were 3’-radiolabeled by using T4 polynucleotide kinase to label cytidine 3’-monophosphate with [γ−^32^P] ATP followed by ligation to the 3’ end of the crRNA with T4 RNA ligase. Radiolabeled oligonucleotides were purified by polyacrylamide gel electrophoresis, eluted in TE buffer (10 mM Tris-Cl, pH 8.0, 1 mM EDTA), and stored at –20°C. Pre-crRNA cleavage reactions were performed using various Cas12a concentrations in 50 mM Na-MOPS, pH 7.0, 120 mM NaCl, 5 mM MgCl_2_, 2 mM DTT, and 0.2 mg/ml BSA and were initiated of a trace amount of labeled pre-crRNA. At various times, 2 μl samples were quenched in 4 μl of denaturing quench (20 mM EDTA, 90% formamide, 0.01% bromophenol blue, and 0.04% xylene cyanol). Samples were analyzed by denaturing PAGE (20% acrylamide, 7 M urea). Gels were exposed to a phosphorimager screen overnight, scanned using a Typhoon FLA 9500 (GE Healthcare), and quantified using ImageQuant 5.2 (GE Healthcare).

### Measurement of pre-crRNA binding kinetics to Cas12a

Pre-crRNA binding was initiated by the addition of a trace amount of radiolabeled pre-crRNA to various concentrations of Cas12a (0.5-10 nM). Reactions were performed in 50 mM Na-MOPS, pH 7.0, 120 mM NaCl, 5 mM MgCl_2_, 2 mM DTT and 0.2 mg/ml BSA as described for the cleavage reactions. At various time points, 2 μl aliquots were withdrawn and added to 4 μl of ice-cold chase solution (reaction conditions plus 100 nM unlabeled pre-crRNA, 15% glycerol, and 0.01% xylene cyanol) and placed on ice. Control reactions in which the chase pre-crRNA and labeled pre-crRNA were mixed and then added to Cas12a ensured that the chase solution adequately competed effectively against the labeled pre-crRNA. After being added to the chase solution, aliquots were placed on ice and then loaded on a 15% native gel run at 4 ^°^C. Gels were exposed overnight and analyzed as described above. Binding reactions in the absence of Mg^2+^ were performed as above but with 5 mM EDTA and no Mg^2+^. Values of *k*_on_ were determined from the slope of the dependence of the observed rate constant on Cas12a concentration.

### Measurement of pre-crRNA dissociation kinetics to Cas12a

For the determination of the dissociation rate constant, we used a 5’-radiolabeled pre-crRNA with deoxyuridine at position −19 to prevent Cas12a-mediated cleavage. Cas12a was mixed with a trace amount of substrate for 1 hr at 25 °C to allow complete binding of the pre-crRNA. Dissociation reactions were performed at 10 nM Cas12a and initiated by addition of the unlabeled pre-crRNA in 50-fold excess over the protein concentration. Aliquots (2 μl) were withdrawn over ∼24 hr, mixed with 4 μl of chase quench (reaction conditions with 15% glycerol, 0.01% xylene cyanol), and stored at 4 °C. Samples were electrophoresed on a 15% native gel at 4 °C, imaged with a phosphorimager, and analyzed as described above. Experiments were performed at 5 mM Mg^2+^ or in the absence of Mg^2+^. For experiments in the absence of Mg^2+^, Cas12a was assembled with the pre-crRNA as described above, and then Mg^2+^ was chelated by addition of 10 mM EDTA.

### Quantification and Statistical Analysis

All binding, dissociation, and cleavage reactions were performed at least three times at the indicated enzyme concentration. Dissociation rate constants are shown as the mean ± SEM of the time courses fit by an exponential decrease. Rate constants for binding and cleavage are shown as the mean ± SEM of the hyperbolic fit of the observed rate constants as a function of Cas12a concentration. All gel-based assays were analyzed using ImageQuant 5.2 (GE Healthcare) and all data fitting was performed using Kaleidagraph.

## ACKNOWLEDGEMENTS

The work was supported by NIH grant R35 GM131777 (to R.R.). S.S. was supported in part by fellowship from the Institute for International Education’s Scholar Rescue Fund (IIE-SRF).

